# Molecular dynamics simulations disclose early stages of the photo-activation of cryptochrome 4

**DOI:** 10.1101/324962

**Authors:** D. R. Kattnig, C. Nielsen, I. A. Solov’yov

## Abstract

Birds appear to be equipped with a light-dependent, radical-pair-based magnetic compass that relies on truly quantum processes. While the identity of the sensory protein has remained speculative, cryptochrome 4 has recently been identified as the most auspicious candidate. Here, we report on allatom molecular dynamics (MD) simulations addressing the structural reorganisations that accompany the photoreduction of the flavin cofactor in a model of the European robin cryptochrome 4 (ErCry4). Extensive MD simulations reveal that the photo-activation of ErCry4 induces large-scale conformational changes on short (hundreds of nanoseconds) timescales. Specifically, the photo-reduction is accompanied with the release of the C-terminal tail, structural rearrangements in the vicinity of the FAD-binding site, and the noteworthy formation of an α-helical segment at the N-terminal part. Some of these rearrangements appear to expose potential phosphorylation sites. We describe the conformational dynamics of the protein using a graph-based approach that is informed by the adjacency of residues and the correlation of their local motions. This approach reveals densely coupled reorganisation communities, which facilitate an efficient signal transduction due to a high density of hubs. These communities are interconnected by a small number of highly important residues characterized by high betweenness centrality. The network approach clearly identifies the sites restructuring upon photoactivation, which appear as protrusions or delicate bridges in the reorganisation network. We also find that, unlike in the homologous cryptochrome from *D. melanogaster*, the release of the C-terminal domain does not appear to be correlated with the transposition of a histidine residue close to the FAD cofactor.

## Introduction

Magnetoreception, the remarkable trait of perceiving Earth′s weak magnetic field, is widespread among the animal kingdom [1]. Yet, the underlying sensory mechanisms have remained elusive. In several instances, most notably the compass sense in migratory birds [2–4], the accumulated evidence supports Schulten′s revolutionary hypothesis of a radical pair-based sensor [5]. According to this model, magnetoreception is the result of the quantum coherent evolution of the singlet and triplet states of transient pairs of radicals under the influence of spin-selective reactions and magnetic interactions [6]. Ritz, Adem and Schulten were the first to speculate that the underlying radical pair could be formed by photo-excitation in the animals′ eyes in cryptochromes [7], a class of blue-light sensitive flavo-proteins that shares some similarity with photolyases [8]; this supposition still applies; to date, cryptochromes are the only vertebrate proteins known to form radical pairs in a physiologically significant photoreaction and their relevancy to magnetoreception has indeed been implicated in a multitude of studies [9–24]. The reader is referred to the many reviews on this subject for a more detailed exposition [6, 23, 25–26].

Four different cryptochromes have been found in the retinae of birds. Among these, two (Cry1a from the garden warbler and Cry4 in chicken) have been shown to form long-lived flavo-semiquinone radicals under photoexcitation [27–28]. Cry1 has been located at the inner and/or outer segments of UV-sensitive cone cells [29–30]. Cry1b was found in the photoreceptor inner segment and the cytosol of ganglion cells [20, 31]. Recently, Günther et al. and Pinzon-Rodriguez et al. have provided circumstantial support for the involvement of Cry4 in magnetoreception [16–17]: According to [16], Cry4 is localized in double cones and long wavelength single cones in the retinae of European robins and chickens. It shows only a weak circadian oscillation [16–17], but its expression is upregulated during the migratory season in European robins [16]. Importantly, as a Type IV cryptochrome, it is able to securely bind the photoactive flavin cofactor, which has previously been questioned for Type II animal cryptochromes including Cry1a from birds [32–33].

The structure of none of the four avian cryptochromes has so far been resolved. A homology model of Cry4 from the European Robin has recently been established [16]. Its three-dimensional structure and folding pattern resembles that of resolved cryptochromes and photolyases and is shown in Figure 1. Its fold is characterized by an α/β-domain and a helical domain. The C-terminal tail (CTT) is thought to mediate signalling functions [28, 34]. In the dark, Cry4 binds a fully oxidized flavin adenine dinucleotide (FAD) chromophore in a central groove [16, 33]. Based on the nomenclature established for the cryptochrome of *D. melanogaster* (DmCry), the structural elements surrounding the FAD-binding site (see Figure 1a) are referred to as the C-terminal lid (residues 420-446 in DmCry; 400 − 421 in ErCry4), the protrusion motif (residues 288 − 306 in DmCry; 273 − 282 in ErCry4), the phosphate-binding loop (residues 249-263 in DmCry; 231 - 248 in ErCry4), and the CTT base loop (residues 154-160 in DmCry; 147 - 151 in ErCry4). Like DmCry and other animal cryptochromes, the protein features an electron transfer chain comprising four tryptophan residues (labelled W_A_, W_B_, W_C_ and W_D_), the tryptophan tetrad [16]. Photo-excitation is accompanied by the reduction of the flavin cofactor yielding FAD^•−^ in the first place. In chickens, this radical can be further photo-reduced to FADH^−^ in a process that likely involves intermittent FADH^•^[28]. It is currently unknown whether the direct photo-reduction of FAD or the dark-state re-oxidation of the fully reduced flavin, FADH^−^, gives rise to the magnetosensitive radical pair underpinning the compass sensor [10, 12, 25, 35]. Some evidence in favour of the reoxidation hypothesis has become available recently [14, 22, 36–41]. Furthermore, a suggestion of how to overcome the issue of fast spin relaxation in the superoxide radical anion, which is closely linked to the latter hypothesis, has been provided [42–43]. In any case, here we focus on the photo-reduction, because it entails a well-defined initial state in terms of a radical pair comprising the flavin anion radical, FAD^•−^, and the radical ion W_D_^•+^. Note, however, that for DmCry comparable structural changes are induced by the photo-reduction and the chemical reduction, suggesting that the charge on W_D_ might not be essential for the induced rearrangements [44].

**Figure 1:**
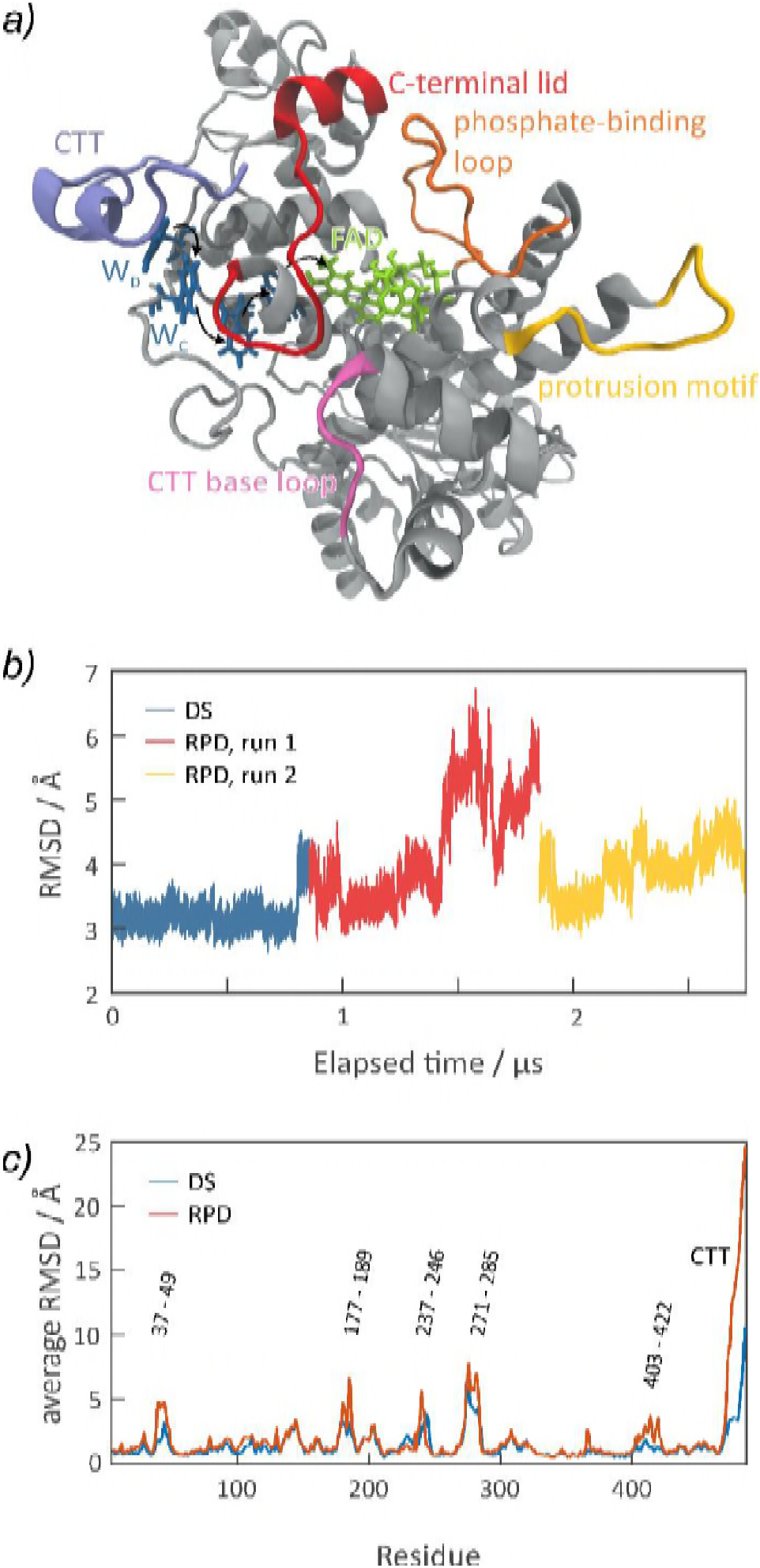
a) Secondary structure model of ErCry4 highlighting the FAD co-factor (lime), the tryptophan tetrad (blue) and pertinent structural motifs such as the C-terminal tail (CTT), the C-terminal lid (red), the phosphate-binding loop (orange), the protrusion motif (yellow) and the CTT base loop (purple). A representative structure of the resting state of the protein is shown, which has been determined through principal component analysis of extensive MD trajectories as part of this study (*vide infra*; R in Figure 3). b) Root-mean-square deviation of the atomic positions of the protein backbone as a function of time for the dark-state (blue) and two independent radical pair-state trajectories (red and yellow). The RMSD has been calculated by aligning the rigid residues to the reference structure from [16]. c) Average RMSD per residue for the dark state and the radical pair-state relative to the representative dark-state structure.

In order to function as magnetic sensor, the radical pair in the cryptochrome must engage in at least two reactive pathways: spin-selective charge recombination, which discriminates the different spin states of the pair, and the competing formation of a signalling state, which induces a cascade of (currently unknown) events eventually giving rise to nervous excitation. In birds, except for the likely involvement of the C-terminal tail, little is known about the conformational changes of the protein related to signalling [28, 34, 37–38]. More specifically, for the cryptochrome 4 from chicken, the molecular accessibility of the C-terminal region to a specific antibody was found to be reduced upon the photo-induced formation of the flavin semiquinone [28, 34]. Light-dependent changes in its trypsin digestion pattern furthermore suggest that the C-terminal tail may bind the FAD-binding domain in the photo-activated state [28]. This suggestion is in stark contrast to the photo-activation of DmCry and the cryptochrome from *A. thaliana* (AtCry), which have been studied in more detail, as well as for the avian Cry1a [37–38]. For DmCry, the reduction of the flavin to the anionic semiquinone by light or chemicals induces a complex conformational reorganisation of the protein, which involves the release of the CTT in addition to rearrangements of its surroundings (the CTT-coupled motif) and several more remote sites [44]. As a consequence, the binding to at least one region of the clock protein Timeless is promoted. These conformational changes were found to be fully reversible under regeneration of the resting state. Based on all-atom molecular dynamics simulations, Ganguly et al. [45] have suggested that the release of the CTT is correlated to a transposition of His378 in the vicinity of the flavin cofactor. The authors furthermore argue that protonation of this histidine could enhance the conformational flexibility of the CTT. In plant cryptochromes, the formation of the anionic radical initiates rapid conformational changes in the photolyase homology region (such as the loss of β-sheet structure in the α/β region), which precedes the slower release of the CTT [46–49]. Here, the anionic flavin radical is eventually protonated by a nearby aspartic acid (D396 in Cry1). The photocycle and signalling mechanisms of plant cryptochromes have recently been reviewed in [50] and the reader is referred to this publication for a detailed account on the topic.

The present study aims to elucidate the early structural changes in the photolyase homology region that accompany the photo-activation of the flavin in a model of Cry4 from the European robin. To this end, we have run extensive molecular dynamics (MD) simulations, which reveal the fast initial response to the photo-induced charge separation on the timescale of a microsecond. We here interpret these data in terms of topological models derived from graph representations of the protein. In addition to the now popular contact maps, we advocate a graph representation that is simultaneously informed by the correlation of displacements in addition to the usual spatial adjacency. We argue that this representation reveals efficient reorganisation networks, which could mediate the signalling in a more directed, purposeful way than networks predicted based on the basis of contact maps. Our approach builds upon two pillars: the use of covariance matrices to analyse the correlated dynamics of protein residues from molecular dynamics, which has for example been described by Hünenberger and co-workers [51] as well as Karplus and co-workers [52–53], and network representations of the protein structure [54] and dynamics [55–56]. In terms of the graph theoretical analysis, our approach is similar to that of Kasahara et al. [55].

## Methods

### Molecular dynamics simulations

All MD simulations were performed using the NAMD program [57] in combination with the CHARMM36 force field including the CMAP corrections for proteins [58–60] and earlier parameterizations for FAD, FAD^•−^ and W^•+^ [61–63]. An equilibrated molecular dynamic ensemble of the dark state (DS) configuration (containing fully oxidized FAD and the standard tryptophan residue) was available from a previous study [16]. The existing trajectory was extended by additional 250 ns following the protocol as outlined in [16]. The last 0.9 μs of the combined trajectory were used in the analysis described here. Two independent MD runs of the radical pair configuration (containing the anion semiquinone FAD^•–^ and an oxidized tryptophan residue at the terminal position of the tryptophan tetrad, W_D_^•+^) were generated from the final dark-state geometry by propagating this state by an additional 1 ns and 2 ns, respectively, followed by swapping the force field parameters for those reflecting the radical pair state (henceforth referred to as RPD). A total of 1.9 μs of MD trajectories was then accumulated for the radical pair state (two independent trajectories; 1 μs + 0.9 μs). All production simulations pertained to the NVT-ensemble; equilibration runs used the NPT-ensemble. A time step of 2 fs was used and temperature and pressure were kept constant at 315 K and, for the equilibration runs, 1 atm using a modified Nosé–Hoover scheme in conjunction with Langevin dynamics. The SHAKE algorithm [37] was used to constrain bonds involving hydrogen atoms to their respective equilibrium distances. A tetragonal box of approximately 109 × 85 × 85 Å^3^ contained 23658 TIP3P water molecules, 74 Na^+^ and 66 Cl^−^ ions in addition to the protein. This gave rise to an overall neutral system at a salt concentration of 50 mM. Periodic boundary conditions were employed to replicate the solvated box in all three dimensions. The particle-mesh Ewald (PME) summation method with a cut-off of 12 Å was used to evaluate Coulomb forces. The van der Waals energy was calculated using a smooth cut-off of 12 Å and a switching distance of 10 Å. The simulation conditions used for the production runs are summarized in Table S1.

### Contact networks / distance maps

In order to account for the different interaction sites of the cofactor, FAD/FAD^•−^ was represented by 5 sub-residues corresponding to the flavin, the ribityl chain, the diphosphate, the ribose and the adenine, respectively (see Figure S1 in the SI). The distance matrices of the centres of mass of all residues and cofactor fragments, **d**(*s*) where *s* ∈{*DS*, *RPD*}, were evaluated for the equilibrated dark-state and the radical pair trajectories, respectively:

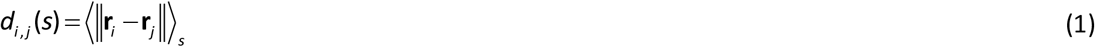

Here, **r**_i_ is the centre-of-mass coordinate of fragment *i* and 〈 〉_*s*_ denotes the average over the DS or RPD trajectory when calculating **d**(DS) or **d**(RPD), respectively. For the radical pair states, the first 200 ns after the charge-shift mimicking the electron transfer were excluded from this analysis.

The protein 3D structural information was mapped onto a graph in which nodes represent residues/sites and edges indicate the existence of neighbourhood relationships. We considered two fragments to be adjacent, i.e. neighbours, if the average distance of their centres of mass was smaller than 8 Å. Subsequently, topological relationships between residues were explored by graph-theoretical tools such as the **geodesic distance** (the number of edges in a shortest path connecting two nodes), **vertex degree** (number of edges incident to a vertex, i.e. number of adjacent residues), the **local clustering coefficient** (a measure of the link-density in the node′s neighbourhood) and the **betweenness centralities** (a measure of the importance of a node defined as the number of shortest paths that pass through a vertex relative to the total number of shortest paths). Following [64], nodes with a degree exceeding 3/2 times the average degree were classified as hubs. For a comprehensive introduction to network representations of protein structures, the inclined reader is referred to [54].

### Displacement correlated contact networks

Global rotational and translational motions of the entire protein were removed by aligning the protein backbone to the average dark-state geometry. The average structure was determined using an iterative scheme: All snapshots were aligned to a template, i.e. the current approximation of the average structure. A new approximation of the average was determined from the aligned structures. The process was repeated until self-consistency of the average was realized. An arbitrary, i.e. the first, structure from the ensemble was used to initiate the method. Following [45], in the molecular alignment step only residues with a root mean square fluctuation of less than 1.3 Å were considered. The RPD structures were aligned to the average dark state geometry using the same set of rigid residues. The protein structure was divided into *N* fragments, which formed the basis for the subsequent analysis. Here, we used each residue of the protein as a fragment, while the FAD cofactor has been represented by five effective residue fragments as discussed above. Let us denote the *i*th component (*i* ∈ {*x*,*y*,*z*}) of the centre-of-mass coordinate of the *k*th fragment by *r*_k,i_ and the 3*N*-dimensional vector of all residue coordinates by **R**. Furthermore, let us refer to the 3-vectors corresponding to the coordinate of fragment *k* as **r**_k_. Thus, **R** is the vector comprising all **r**_k_ 3-vectors as partitions, i.e. **r**_k_ = (***R***_3k−2_, ***R***_3k−1_, ***R***_3k_). The 3*N*-dimensional covariance matrix **C** was computed from

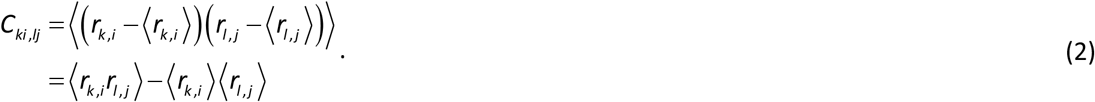

Here, the brackets 〈 〉 signify the average over the MD trajectory. **C** as defined in eq. (2) should be rescaled as **C**′ = *M*/(*M*−1) **C**, with *M* denoting the number of averaged structures, in order to arrive at the best unbiased estimate of the covariance matrix if the observations are from a normal distribution, as would be expected for structures subjected to thermal motion (here *M* = 95,700 and 189,300 for the Ds and RPD state, respectively). The 3*N* orthonormal eigenmodes of the *N* fragments, **Q**_k_, were determined as the solution of the eigenvalue problem

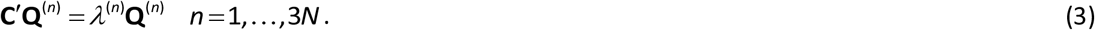

By partitioning **Q**^(n)^ in the same way as **R**, it is possible to associate 3-vectors 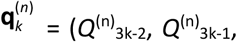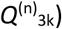) with every fragment *k*. The displacement **R**−〈**R**〉 can be re-expressing in terms of the eigenmodes by evaluating:

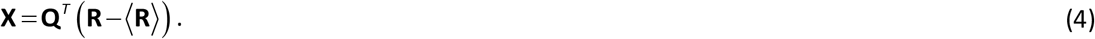

The variance of the *k*^th^ 3-vector in **X**, **x**_k_, due to the *n*^th^ normal mode is then given by 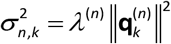. The total variance in **x**_k_ is 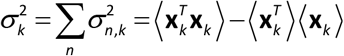.

The following heuristics were used to identify correlated motions of fragments. A simple approach can be directly based on the correlation matrix **P**, which is related to **C**′ by

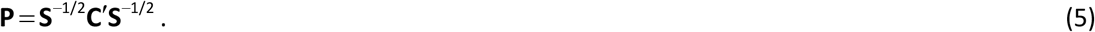

Here, **S** is the diagonal matrix comprising the variances of the displacements of the individual fragments, i.e. the matrix assembled from the diagonal of **C**′. Fragments *k* and *l* have been classified as correlated if the correlation coefficient of any pair of coordinates exceeds a problem specific threshold, *r*_min_. In detail, denoting the associated *N*-dimensional adjacency matrix by **M**, we may define [55]

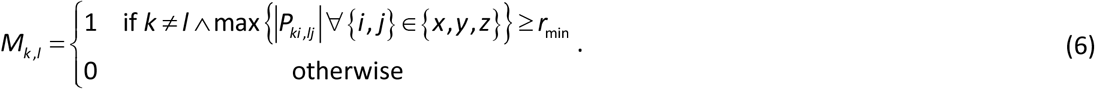

An alternative approach to detect correlated reorientational motions of fragments is based on the eigenmode decomposition, eq. (3). Following [65], two fragments *k* and *l* are considered correlated if the same normal modes provide, on average, similar contributions, i.e. if the ordered sets of displacement contributions 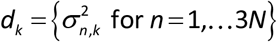 of the two fragments are positively correlated. The degree of correlation of *d*_k_ and *d*_l_ can be expressed by the Pearson correlation coefficient, corr(*d*_k_, *d*_l_). Again, one has to binarise the correlation measure with a suitable *r*_min_′ to obtain the adjacency matrix of motional correlation **M**′

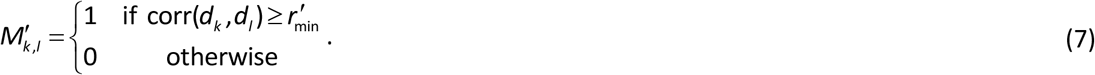

In practice, it is advisable to choose the cut-off distances in eq. (6) or (7), *r*_min_ or *r*_min_′, to be as large as possible while ensuring that the resulting graph comprised a single dominant connected component in addition to a few isolated fragments. For comparable cut-offs the adjacency matrix **M**′ tends to be less sparse than **M** and comprise more large-distance correlations. In combination with the additional adjacency criterion (see below), comparable results are obtained from both approaches.

Our analysis aims to identify spatial networks of ErCry4 fragments that provide a pathway for structural rearrangements. In other words, we are interested in networks of local correlations that function as the building blocks of the overall reorganisation dynamics. This view stipulates that not all correlations from **M** are considered, but only those that pertain to neighbouring interacting sites. In order to accommodate this adjacency aspect, we introduced the matrix **D**:

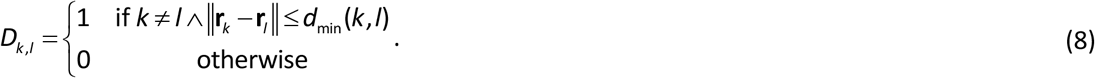

Here, we have used *d*_min_(*k*,*l*) = 8 Å if fragments *k* and *l* are not part of the same rigid secondary structure element. If, on the other hand, *k* and *l* belong to the same α-helix or β-strand (as determined by the STRIDE program [66]), *d*_min_ = ∞; the latter condition reflects that correlated motions over rigid secondary structure elements were always considered as direct link, thereby simplifying the interaction network by providing “shortcuts“. This is consistent with our observation that for the ErCry4 protein considered here, secondary structure elements are preserved and exhibit collective modes.

The displacement correlated contact network graph was eventually generated using the adjacency matrix **A** as defined by

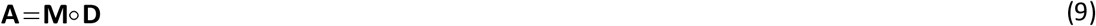

or a similar expression with **M′** replacing **M**. The “∘” denotes the Hadamard entrywise product. This graph was analysed using the same topological measures as set out for the contact networks above.

## Results

### Root-mean-square deviations, solvent accessible surface areas and interaction energies

Figures 1b and S2 in the SI illustrate the temporal evolution of the root-mean-square deviation (RMSD) of the position of the backbone atoms (C, O, N, CA) in ErCry4 with respect to the reference structure from [16]. It is immediately obvious that the charge shift that generates the radical pair state (RPD) from the ground state molecule (DS), gives way too large fluctuations and an increase of the RMSD. The mean RMSD per residue (Figure 1c) reveals that large RMSDs are accrued at the CTT and, to a lesser extent, several loop regions (which are to be specified in detail below). Note that in general sites of large RMSDs coincide for the DS and RPD state. However, for the latter the RMSD are typically bigger. The largest differences of mean RMSD per residue for the DS and RPD state are observed for the CTT and the mobile regions surrounding residues 45, 185 and the region following the C-terminal lid around residue number 410. While the RMSD plots reveal interesting features, this analysis is biased by the choice of the reference structure and the protocol for structure alignment. These limitations can be overcome by comparing the average DS structure to the average RPD structure of ErCry4 in internal coordinates (see below).

The photo-activation is accompanied by a substantial increase of the solvent accessibility of W_D_; its solvent-accessible surface area (SASA) increases from 26.5 Å^2^ to about 72 Å^2^, as is shown in Figure S3 in the SI. Likewise, the SASA is slightly larger for FAD^•−^ in the RPD state than FAD in the DS (26.5 Å^2^ vs. 30.3 Å^2^; see Figure S4). The Coulomb interaction energy of the FAD cofactor and W_D_ increases from −0.31 ± 0.18 kcal/mol in the DS to −10.64 ± 0.68 kcal/mol in the RPD state (see Figure S5); the standard deviation increases by a factor of 3.7. As a consequence of the additional charge, the binding energy of FAD is furthermore more negative in the RPD state than the DS; the standard deviations of the binding energy are comparable in both states (Figure S6).

### Distance matrices and contact maps

In order to assess the internal structure of ErCry4 and its changes upon photo-ionisation, we have evaluated the average distances of the centres of mass of all residues over the equilibrated trajectories. Unlike many other studies that employ the distances of C_α_-atoms, we have focused on centres of mass as their location is more sensitive to sidechain reorganisations. The cofactor, FAD, has been represented by five effective residues corresponding to the flavin, the ribityl chain, the diphosphate, the ribose and the adenine, respectively, as explained above and illustrated in Figure S1 in the SI. The difference of the average distance matrices of the RPD and DS ensembles, eq. (1), reveals the identity of segments that undergo major restructuring upon activation. Figure 2 provides a density plot of this difference metric. It exhibits a checkered pattern, which is indicative of mobile segments that change their relative orientation with respect to a rigid mainstay structure. The total absolute displacement shown in Figure 2 is defined as

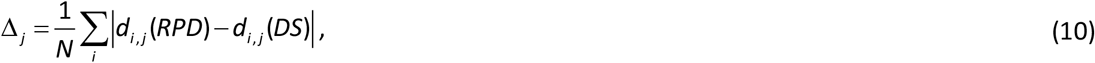

with *d*_i,j_ denoting the distance of the centres of mass of fragments *i* and *j*. This quantity reveals the identity of the mobile residues (top panel in Figure 2). The dominant structural changes occur in the C-terminal region, in particular the segment comprising residues 472 to 486, which moves away from the core of the protein while approaching the distal α-helix H17 (residues 450 to 456). The conformational changes are, however, not limited to the C-terminus. The difference metric Δ_j_ reveals additional reorganisation of the helix-bound residues Arg419 and Thr420 and several surface-exposed loop regions: residues 405 – 415, 276 – 285 (which correspond to the protrusion motif), 237 – 246 (which covers most of the phosphate binding loop), 179 - 186 and 38 – 48. In addition, large shifts of a few solitary residues are observed (e.g. Arg209, Asp367 and Ser130). We have also constructed contact maps (adjacency matrices) by binarizing the average distance matrices with a cut-off distance of 8 Å. The difference of the contact maps of the RPD and DS is indicative of the reorganisation process giving rise to the formation of new, or the interruption of existing, interactions (see Figure S7 in the SI).

**Figure 2:**
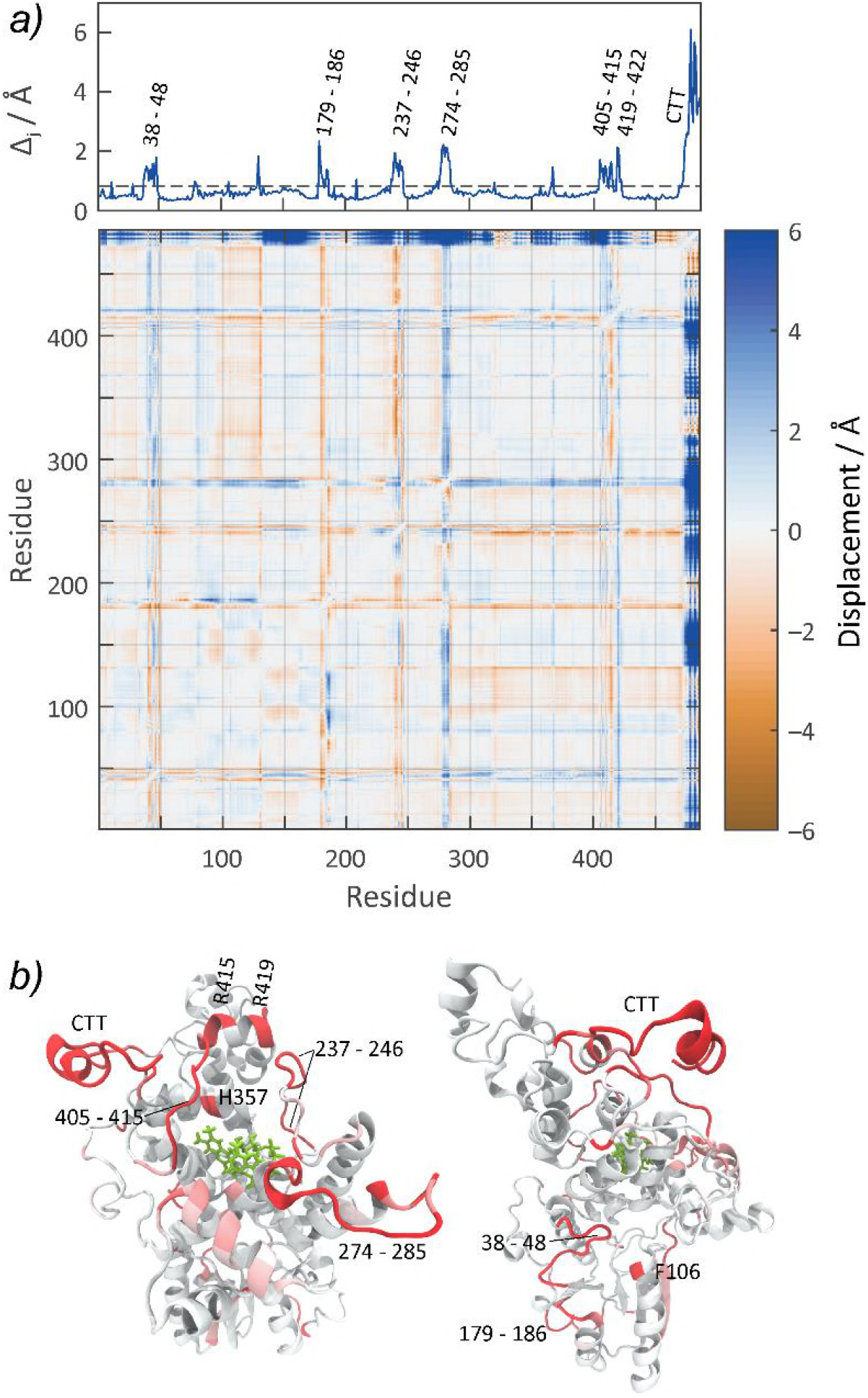
a) Differences in the distances of the centres of mass of the residues of the DS and the RPD states of ErCry4. The top panel gives the absolute relative displacement as defined in eq. (10). For mobile sites (Δ_j_ *N* > 400 Å; see eq. (10); the dashed line indicates the threshold value), the sequence numbers of the affected residues are indicated. b) Illustration of the secondary structure of ErCry4 with mobile sites indicated in red.

While the combined analysis of the distance matrices and the contact maps cannot reveal the temporal/causal chain of events that facilitate these reorganisations in ErCry4, the correlated transpositions of a few key residues appear noteworthy. For example, the restructuring of the loop 237 – 246 appears to correlate with a reduced distance between Ser245 (in the loop) and Lys234 (change in average distance: −3.4 Å). The latter directly interacts with the diphosphate group of FAD via an H-bond involving its ammonium group. Likewise, Arg209, one of the isolated residues that are strongly displaced, decreases its distance to Pro244 and increases its distance to the rest of the loop. Figure S7 in the SI reveals that the reorganization of the loop is accompanied by the formation of a series of new contacts with residues 337 – 350, a mostly α-helical segment that itself does not undergo strong reorganisation, and the segment 417 – 430. In terms of the difference metric, this is evinced in the displacement of the adjacent residues Arg419 and Thr420 and its upstream segment 405 – 414. While the former appears to be transpositioned towards the C-terminal domain of ErCry4, the upstream segment 405 – 414 mostly shifts in the N-terminal direction. In doing so, residues 413 and 414 establish new contacts with an adjacent α-helix (around residue 357). This is one of two reorganisation motifs in the vicinity of the C-terminal segment. The second involves repositioning of Asp367, which is located close to the oxidized W_D_ = Trp369. Unlike Asp367, W_D_ does not strongly change its relative position with respect to the rest of the protein upon photo-ionisation.

The reorganisation of ErCry4 is not confined to the C-terminal part of the protein, but extends to the loop region 276 – 285 and the N-terminal part. Large displacements can be observed for residues 38 – 48, which correlate with a reduced Lys377-Ser45 distance. The restructuring of this loop goes along with the formation of a new contact of Arg39 and Asp197, which likely conveys additional rearrangements in the adjacent loop covering residues 179 – 186. Here, the largest displacement is observed for the surface-exposed cysteine Cys179. Large displacements are also detected for residues 276 – 285, the protrusion motif; a surface exposed loop adjacent to one of the α-helices forming the FAD binding pocked.

The residues surrounding the flavin cofactor are comparably immobile. The largest total displacement, Δ_*j*_, of a residue within 15 Å of the isoalloxazine ring is due to His357, which is displaced by 0.6 Å away from the flavin; see Figure 2. This is remarkable because a similar displacement of a solitary histidine has been implicated in the signalling of DmCry [45]. While these interaction motifs do indeed appear comparable on the first glance, it here involves a different residue. Based on a sequence alignment of DmCry and ErCry4, the histidine that was expected to mediate this mechanism in ErCry4 was His353 (corresponding to His378 in DmCry), which, however, exhibits an insignificant total displacement, Δ_j_, and a smaller shift with respect to the isoalloxazine (−0.3 Å). Furthermore, His357 makes a new contact with Phe411 and approaches the related loop. The largest individual shift of close-by residues with respect to the isoalloxazine is due to Leu383 (0.9 Å).

### Principal component analysis

A principal component analysis (PCA) on the motions of the centres of mass of the ErCry4 residues was carried out following the procedure outlined in the Methods section (for additional information, see [67]). The combined RPD and DS trajectories were analysed simultaneously with the aim of identifying the predominant reorganisation modes associated with the photo-induced charge separation and their large-scale correlations in ErCry4. A small number of principal components (PC) was found to account for the observed structural variability. The dominant PC explained 33 % and the three leading PCs 64 % of the total variance (relative contributions calculated as *λ*^(*i*)^ /Σ_*j*_^*λ*(*j*)^; cf. eq. (3)). The corresponding biplots are shown in Figure 3. These plots provided the probability densities for the system configurations expressed as a function of their projection onto the normal modes (cf. eq. (4)).

Several characteristic conformers can be identified from the biplots. Three conformers have been marked and their 3D-structures plotted in Figure 3. Conformation R is representative for most of the DS-trajectory; A1 and A2 correspond to two conformers occurring predominantly for the photo-activated state. The two main conformers, R and A1, are separated by a translation along PC1 for constant PC2. Note however, that the direct reorganisation path appears to involve a substantial free energy barrier, which the system can avoid by paths intermittently changing PC2 and PC3 (and/or lower order PCs; see the depopulated region between R and A1 along the PC1 component in Figure 3). The major reorganisations are linked to the mobile residues that have already been named above based on the analysis of the distance matrices. Figure S8 in the SI shows the weighted RMSD modes [67], which provide the relative contribution of every residue/fragment to the total variance. It also reveals that the reorganisations at different sites are highly correlated, i.e. they proceed in a concerted fashion for changes in a single coordinate. Most mobile sites contribute significantly to all four dominant PCs. The notable exception is the reorganisation of residues 80 – 135, which appears to be an exclusive feature of PC2 (among PC1 to PC4; see Figure S8). Expectedly, the largest per residue contribution of the dominant PCs is due to the CTT. The site surrounding residue 45 accounts for the second largest contribution in PC1 and PC2. The major reorganisations from R to A1, which are predominantly accounted for by PC1, are found at the CTT and the CTT-coupled motifs including the C-terminal lid, the phosphate binding loop and the protrusion motif (see Figure 4). In addition, marked structural changes occur at the α-helix containing Asp197 and its adjacent loops (residues 186 to 206), which increase the solvent exposition of the α-helix. Furthermore, the loop region comprising residues 38 – 48 at the N-terminal end is internalised. For A2, the structural changes follow a similar pattern. However, the CTT is rotated away from the protein surface. Interestingly, for this structure, the loop region at residues 38 – 48 coils to form an α-helix (see Figure 4, left), which is exposed at the protein surface, quite contrary to what is observed for PC1 at this site. Taken together, our analysis suggests that, just like in cryptochromes from other species, the CTT and the CTT-coupled motif show major responses to photo-ionisation. In addition, the loop region 38 – 48 at the N-terminus is identified as a site that undergoes interesting structural transformations upon photo-ionisation.

**Figure 3:**
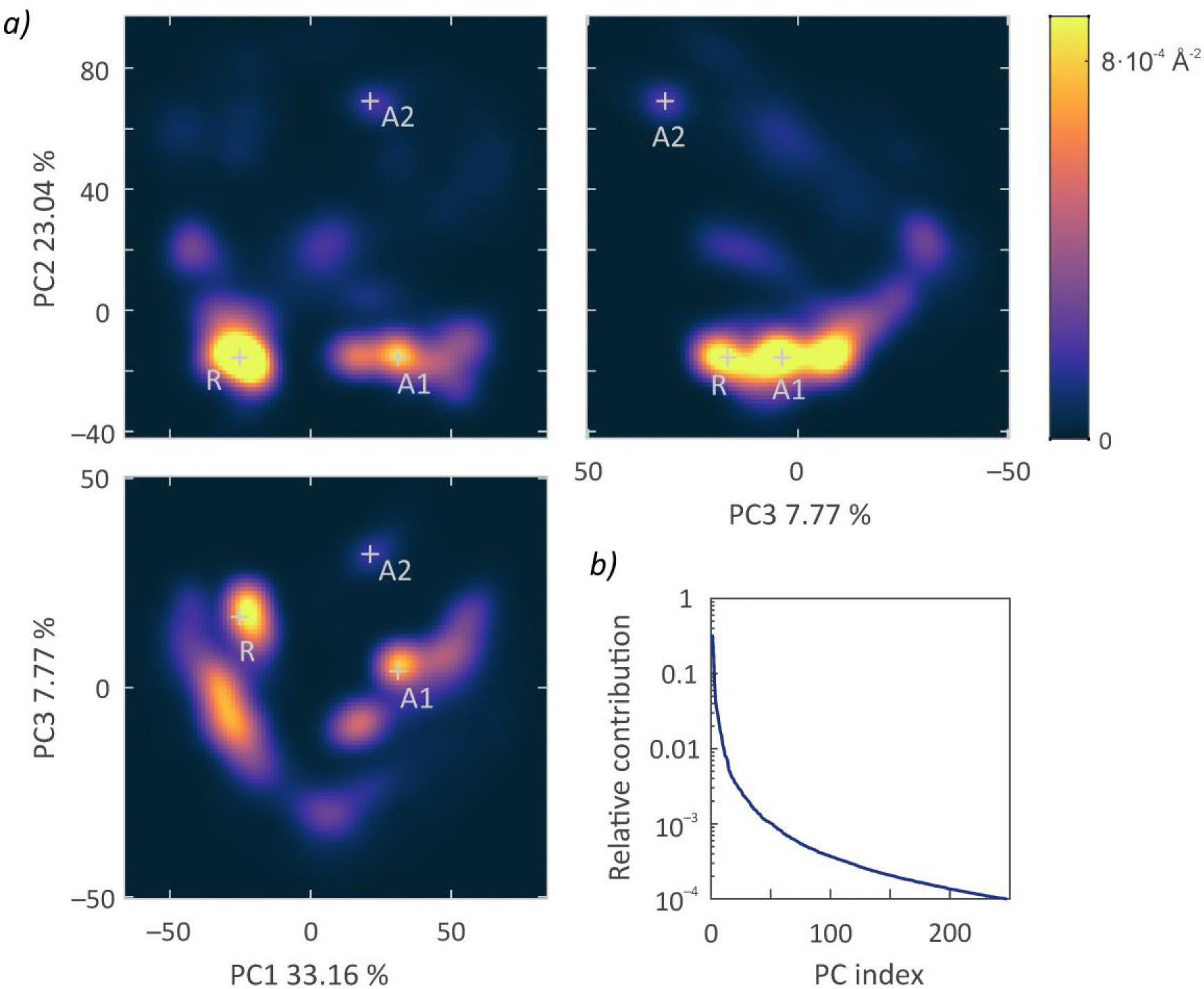
a) Biplots of the configurational ensemble sampled by the DS and RPD states of ErCry4 during the accumulated MD trajectories. The probability density of these configurations is plotted as a function of the displacement along the first three principal component modes, i.e. the dominant collective reorganisation modes. The displacements along the individual modes have been evaluated from eq. (4) and are given in units of Å. The labels specify the relative contributions of the modes to the overall variance. b) Relative contributions of the principal components to the motional variance as a function of their ranked index.

**Figure 4:**
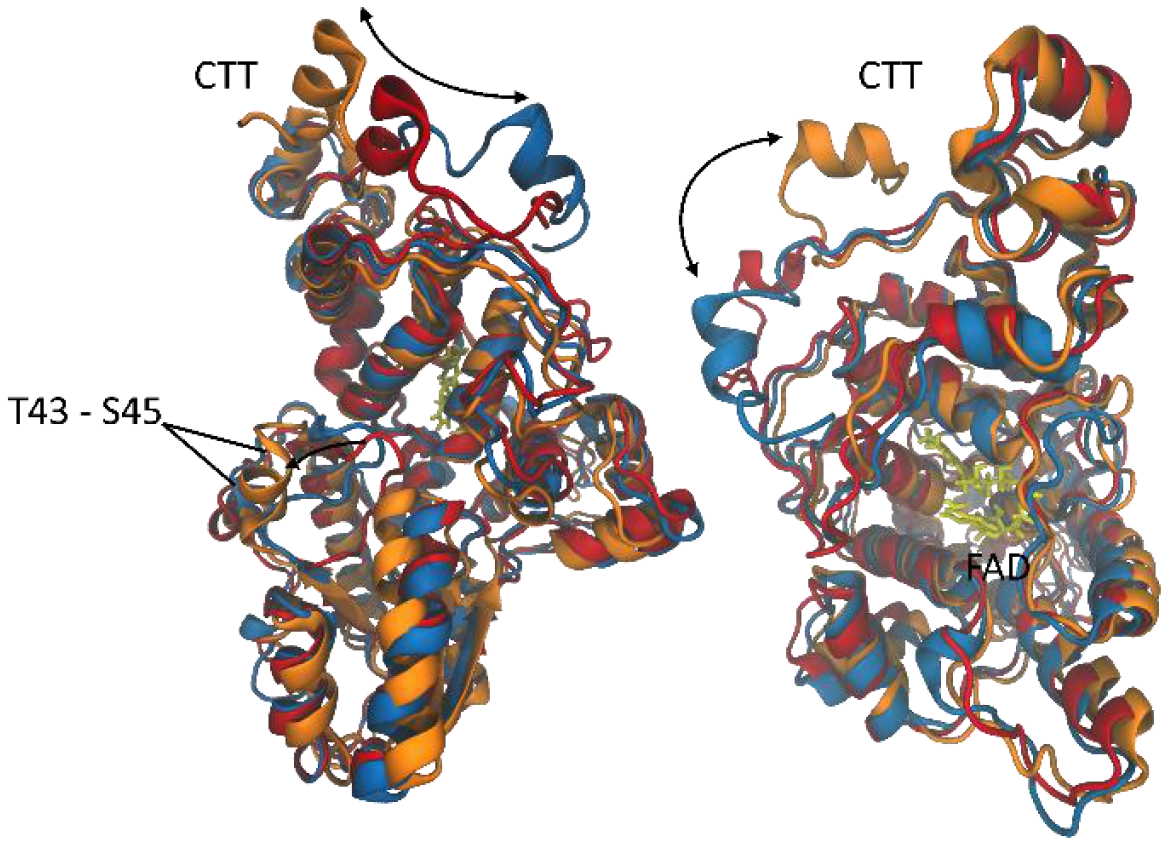
Representative conformers sampled by ErCry4 during the MD trajectories. Blue: resting state conformer (R). Red (A1) and orange (A2): Conformers interpreted as activated states with the CTT released.

### Protein contact networks / adjacency graphs

Contact networks provide an alternative representation of the structural features of proteins. The graph representations were generated from the distance matrices of the DS and the RPD states according to the procedure described above. A graphical representation of the contact network of the DS is shown in Figure 5; the plot has been obtained by using the spring-electrical graph embedding of Wolfram Mathematica [68]. Characteristic nodes have been highlighted by colours; mobile sites for which *N* Δ_j_ > 400 Å (cf. eq. (10)) have been drawn in yellow; the isoalloxazine subunit of the cofactor (henceforth denoted F) is shown as a blue circle; other features will be discussed below. The corresponding graph representation of the RPD state of ErCry4 is shown in Figure S9 of the SI. Some of the mobile sites are obviously associated with peripheral loops or extensions. The CTT part of ErCry4 is particularly prominent (even more so for the RPD state; see Figure S9), as is the loop in in between the helices H6 and H7 at the N-terminal side of the protein and the loop connecting the helices H9 and H10. The prominently reorganising site surrounding residue Ser45 is located at the bottom of the graph, although not at its topological surface. Many mobile residues appear to be scattered over the network without following an obvious pattern. Comparing Figure 5 with its counterpart for the RPD state of ErCry4 in Figure S9 reveals the structural changes associated with the photo-activation. The release of the CTT is striking as are the reorganisations at residues 180 and 278, which appear to assume a more “exposed” position. The mobile residues around Ser45-are moved to the bottom surface of the graph. These changes are in accordance with the picture of an opening of the protein structure, which has also been postulated to accompany the activation of other cryptochromes [44–49].

The notion of an opening of the protein structure is also supported by analysing the geodesic distance, i.e. the number of edges in a shortest path connecting two nodes in the network; see Figure S10 in the SI. In fact, for the DS state of ErCry4, the flavin residue F is classified as a central vertex suggesting that its vertex eccentricity (the largest distance to other vertices) provides information about the protein opening as well as the length of the reorganisation pathway from F to any effector site in the periphery. We find that the eccentricity of F increases from 8 to 11 upon ErCry4 photo-excitation while the graph diameter increases from 15 to 18. We have calculated the difference of graph distance matrices. It reveals distance changes upon photo-activation ranging from −6 (expansion) to +2 (contraction). Among the preserved secondary structure elements, one particularly notes that the helices H7 and H8 approach F, while the helices H3 and H17 are drifting away from F (see Figure S10).

**Figure 5:** Protein contact network representation of the DS state of ErCry4 as derived from the average distance matrix, eq. (1), of a 0.9 μs MD trajectory. Mobile nodes, hubs, and the 20 nodes of highest betweenness centrality have been highlighted in colour (see legend). The fragments contributing to the FAD cofactor are shown in blue and the tryptophan tetrad in light grey. Individual residues and prominent mobile regions have been labelled by their residue sequence numbers.

Figure 5 also features nodes that are deemed important for the structural identity of the network and the transmission of signals. This includes the so-called hubs, nodes with large vertex degree (i.e. the number of edges incident to a vertex; for hubs it is expected to exceed the mean node degree by at least 3/2), and the 20 nodes of highest betweenness centrality, which is defined as

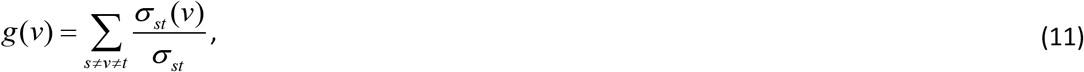

where σ_st_ is the number of shortest paths between nodes *s* and *t* and σ_st_(*v*) is the number of shortest paths that pass through vertex *v*. g(*v*) is a measure of the importance of node *v* to the connectivity of the network. In particular, the presence of high-betweenness nodes has been suggested to facilitate long range transmission of information in protein networks [54]. Here, 5 % of the nodes are classified as hubs, which is a typical fraction for protein networks [64].

As far as our aspiration to identify mobile sites in ErCry4 is concerned, additional insights are provided by the vertex degree, the local clustering coefficient and the node centrality. In fact, sites comprising more than 1 residue have an average degree smaller than the overall mean degree (9.2 and 9.1 for the DS and the RPD state, respectively). Furthermore, for most mobile sites, residues tend to cluster together forming tightly knit groups characterised by a relatively high local clustering coefficient. On the contrary, the betweenness centrality (or likewise the eigenvector centrality) is low for the mobile residues. This portrays the image of mobile sites as groups of peripheral residues that are only weakly coupled to the core protein. Additional details, including representations of the mentioned graph-theoretical parameters as a function of residue sequence number, are provided in the SI (Figures S10 to S12).

### Displacement correlated contact networks

Contact networks provide an accessible way to describe the structure of the DS and RPD states of ErCry4 and their differences. However, it lacks the insight as to how the underlying conformational changes are propagated through the protein, at least beyond the level that can be guessed based on mere adjacency. This is what displacement correlated contact networks seek to deliver by combining the concepts of correlated motion and closeness. Figure 6 shows the cross-correlation network as obtained based on the correlation of the displacements of the centres of mass of the residues over the combined DS and RPD trajectories. The correlation is measured here as expressed by eq. (10) in combination with a threshold correlation coefficient of *r*_min_ = 0.7. The network in Figure 6 has an astoundingly simple structure, for which most mobile sites are easily recognisable either as protrusions (e.g. CTT, protrusion motif, phosphate binding loop) or feeble bridges (e.g. C-terminal lid). These signalling sites are connected to tightly coupled core structures, by but a few residues. The network also shows evidence of dynamical bottlenecks between the left and right side in Figure 6, which roughly separates the N-terminal part from the C-terminal part of the sequence. Not all of the residues are coupled to the main subgraph shown in the figure; a few isolated residues do not show a tendency for correlated motion with neighbouring sites. Among these are the mobile sites Met46, His47, Ile48 and His357. Evidently, this is a function of the cut-off value used in valuing correlation of motion in the protein, which here was chosen to produce a graph as sparse as possible but subject to the constraint that all subgraphs that are uncoupled from the core graph comprise only single residues. This allows the essentials of the correlated restructuring motion to be captured, while keeping the network simple. Here, this procedure gives rise to a core connected component of graph diameter 26 and a vertex eccentricity of F of 14 (as compared to 15 and 8, respectively, for the DS contact network detailed above). The most notable feature of this graph is the high density of hubs, which here account for 20 % of the nodes as compared to 5 % for the contact networks. In addition, the assortativity (the tendency of nodes to attach to nodes of similar degree) of the correlation network is higher than for the adjacency networks. Mobile sites are typically characterised by a lower degree, high local clustering coefficient and low betweenness centrality. In addition, it is noteworthy that all signalling sites except for that surrounding Ser45 are remote from F and thus the network centre. Detailed summaries of the pertinent graph-theoretical parameters are given in the supporting information (Figures S13 and S14); we will return to peculiar features in these data in the Discussion. Expectedly, the displacement correlated contact networks do depend on the choice of the correlation metric and the associated cut-off. While the network layout and individual paths can be strongly affected if different methods retain (slightly) different dynamic correlations, the overarching features such as trends in the pertinent parameters, the identity of important nodes/hubs agree. In favour of conciseness, here we shall limit ourselves on the graph based on eq. (10).

**Figure 6:** Displacement correlated contact network of ErCry4. The graph combines the adjacency information of the ground state (eq. (8)) with the correlation of motion of the combined DS and RPD trajectories (eq. (6), cutoff: *r*_min_ = 0.7) in an attempt to capture the reorganizational dynamics relevant for the activation and signalling of the protein. See the legend of Figure 5 for additional details.

## Discussion

### Structural changes upon photo-activation

Our data suggest that *grosso modo* the photo-activation of ErCry4 is associated with an expansion of the protein. In particular, the CTT is attached in the DS and, when illuminated, released in a fashion similar to what has been described for AtCry and DmCry [44–50]. We do not see indications of the CTT’s attaching to the photolyase homology region (PHR) of the protein in the course of the photoactivation, as suggested for cryptochrome 4 from chicken [28]. In [28], the authors found that the accessibility of trypsin to the FAD-binding domain is reduced upon light illumination, which they interpreted as the binding of the CTT to the protein core, thereby protecting the associated tryptic digestion sites. The present analysis does not support this suggestion. On the contrary, the CTT appears to be released in a way that increases the accessibility at the FAD binding site (see Figure 4). The photoactivation of ErCry4 is, however, linked to substantial reorganisation at the CTT-coupled motif, which could possibly alter the susceptibility to trypsin. Alternatively, the discrepancy could be attributed to differences in the conformational changes of the full-length protein relative to the truncated structure currently amenable to theoretical studies or to the protonation state of the flavin. Here, we have investigated the radical anionic state, as this is the state that, in AtCry and DmCry, accounts for the fast, microsecond, structural rearrangements that lead to protein signalling. Indeed, in AtCry these conformational changes precede the protonation of the flavin anion radical at the nitrogen atom N5 with a nearby aspartic acid functioning as a proton donor (D396 in Cry1) [61–63, 69–72]. As no comparable proton donor exists in ErCry4 (just as in DmCry; see Figure S15 in the SI), one can safely assume that the microsecond conformational dynamics are correctly represented in our calculations. Further studies will be necessary to assess the influence of the full-length CTT once its structure has been revealed or a suitable template for homology modelling established.

### Signalling mediated by mobile histidines

Ganguly et al. [45] have conducted a detailed atomistic study of the activation of DmCry and concluded that the CTT release correlates with the conformation of conserved His378 (DmCry numbering), which resides between the CTT and the flavin cofactor. Given the high degree of homology of DmCry and ErCry4 (45.5% sequence identity over the entire sequence), in particular for the PHR, the question arises if a similar conformational change could be associated with the signalling of ErCry4. His378 in DmCry corresponds to His353 in ErCry4. According to the presented data, His353 in ErCry4 does not undergo a significant shift with respect to the isoalloxazine subunit of FAD (F) upon photo-activation (shift: −0.3 Å, in the opposite direction and by less than observed for His378 in DmCry). Furthermore, His353 assumes a peripheral position in the displacement correlated contact network, which does not hint at an important role in the activation of the protein’s CTT domain (see Figure 6). In particular, the residue does not qualify as hub or important residue from its betweenness centrality. Alternatively, here His357 is realized as a mobile residue in the vicinity of FAD (distance to isoalloxazine ring: ~ 9Å). However, our correlation network analysis indicates that its displacements are only weakly correlated to the mainstay motions, i.e. mostly random, to the extent that the node is not coupled to the main reorganizational network. Thus, it is likewise unlikely that His357 in ErCry4 could perform a similar function to His378 in DmCry. A closer inspection of the protein environment of His353 in ErCry4 reveals that its complacent signalling role does not come as a surprise: Unlike for DmCry, the fold of ErCry4 does not allow hydrogen bonding of the histidine to the ribityl unit in FAD. This applies to all protonation states of His353. Correspondingly, we have not yet explored the impact of prototropy or double-protonation for ErCry4, which are relevant for DmCry [45]. Obviously, this view might have to be altered if the crystal structure, once available, showed a fold of the photolyase homology region differing from the predictions of the homology model used in the present investigation. This cautioning remark aside, note that the model is supposed to be of high quality for this part of the protein [16].

### Signalling pathways

Signalling “pathways” are often understood as isolated continuous chains of residues connecting the key sites of a protein or a group of proteins, i.e. the effector site and possible multiple recipient sites. The present analysis based on networks of short-range correlated displacements portrays an entirely different picture of ErCry4. Instead of single pathways, we have identified difficult-to-define networks of residues that mediate global conformational changes involving most if not the entirety of the protein. Specifically, we have established by PCA, that two main conformers, interpreted as resting and activated, are linked to changes in a single principal component that simultaneously affects multiple sites including the N- and C-terminal parts of the protein. Furthermore, we have realized that the correlated reorganisation network comprises communities of densely coupled nodes with a remarkably high density of hubs (20 % over the entire protein as opposed to 5 % in contact networks of average proteins; even dense in the strongly coupled subdomains). For these regions, the coupling network appears highly redundant and able to transmit information in a concerted fashion. The question of a pathway in the sense of a chain of well-defined residues is obviously moot for such tightly knit sections. On the other hand, many of the mobile sites in ErCry4 are characterized by a linear or quasi-linear topology, for which the notion of a pathway applies in good approximation if the view is limited to the dynamics away from the interconnection site. The mentioned, densely coupled communities can be determined from the graph by maximizing the so-called modularity, i.e. the fraction of the edges that fall within communities minus the expected fraction if edges were distributed at random [54]. This procedure yields communities of dense connections within reorganisation modules but sparse connections to adjacent modules. In Figure 7 we show a 3D representation of ErCry4 for which the 12 communities of cardinality larger than 1 have been coloured differently. According to the above discussion, within these communities signals are transmitted either linearly (for the mobile protrusion of the graph) or in a complex multi-sited mode involving many concurrent pathway. While the simple pathway picture is ill-conceived in the presence of the latter, the reorganisation network is still depending on a few distinguished nodes, the nodes of high centrality. These nodes mediate bottlenecks in the reorganisation network, typically at the interface of the tightly coupled communities. In this way, they facilitate the long range transmission of information while the hubs support short to medium range reorganisation. Note that for ErCry4 all mobile residues but residues 405 – 407 are located in a community other than that containing the flavin cofactor. Consequently, high betweenness nodes are essential for most reorganisations in this protein. In Table 1 we have collected the 20 nodes of highest betweenness centrality as well as the hubs. Some of these nodes are also labelled in Figure 6. The most remarkable feature is the fact that 4 residues (Glu102, Met103, Ser250 and Asp385) are crucial in passing the signal from the F-residue to sites at the N-terminus of the protein; if one wished to abolish this pathway, these would be the residues to target. While high betweenness residues also appear for the connection of F and the CTT domain, the system is more redundant at the C-terminal end suggesting that a point mutation will have a smaller impact. Figure 7b illustrates the coarse grained network graph of reorganizational communities. There, the widths of the edges are proportional to the number of pairs of residues transmitting the coupling. This picture suggests that the FAD-site (community 2) is efficiently coupled to community 1, which mediates the release of the CTT (community 8). The mobile site involving residue 45 (contained in community 4) is directly innervated by the co-factor bearing community 2. Table 2 summarizes the association of mobile sites with the reorganizational communities they are contained in. It is remarkable that only communities 11 + 1 and 4 + 12 appear to be coupled through mobile sites at the interface of the respective pairs. All other links involve high betweenness sites that however are only displaced by a comparably small amount with respect to the protein backbone. Note furthermore that while residues 405-415 are located at the interface of communities 2 and 11, they do not provide a connection via correlated displacements.

**Figure 7:**
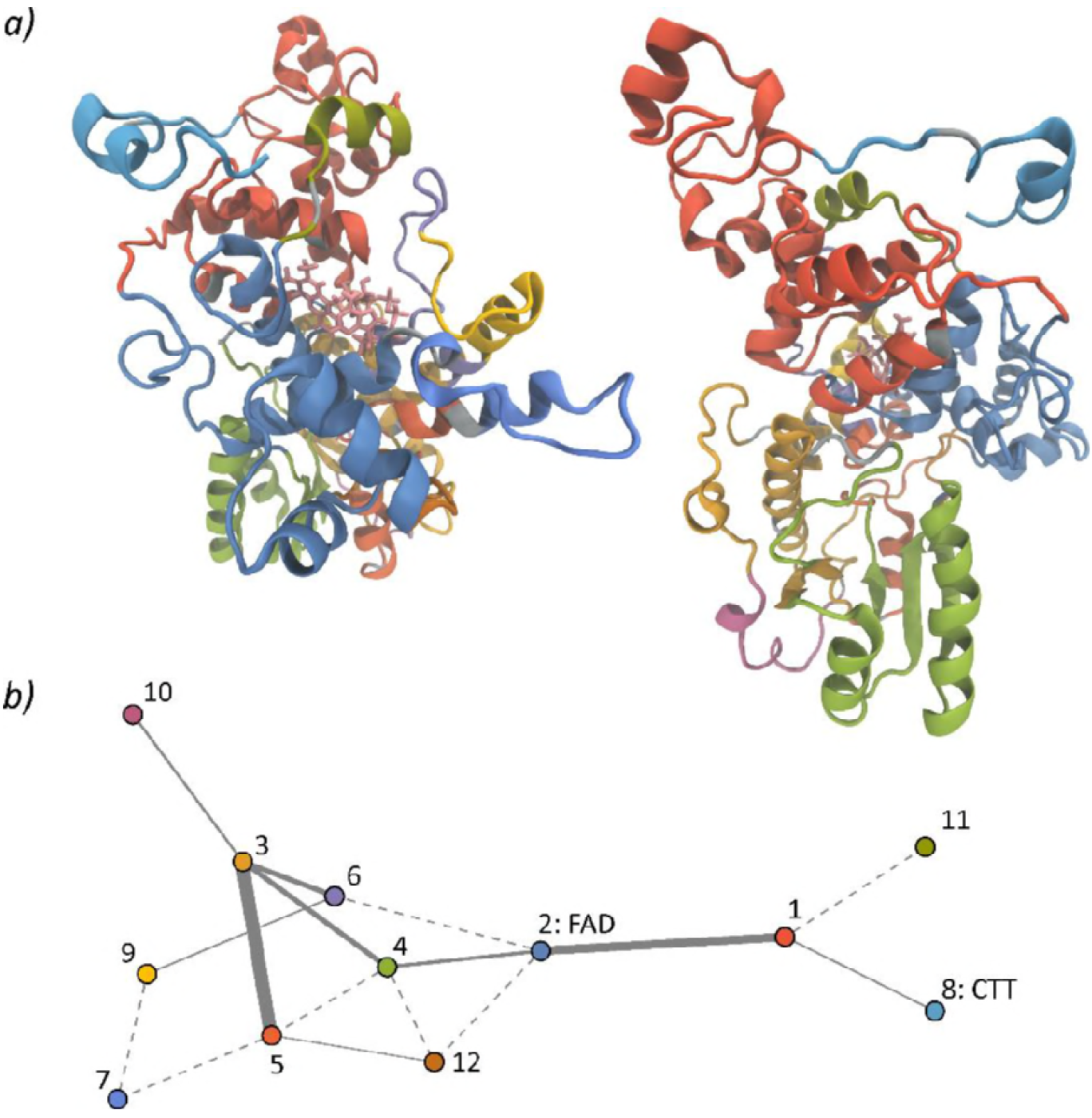
a) Secondary structure of ErCry4 with residues coloured by their associativity to reorganizational communities as derived from the graph in figure 6 by maximising the graph modularity. b) Coarse grained network of reorganizational communities. The colours of the nodes correspond to the colours used for the 3D structures above. The widths of the edges are proportional to the number of connecting pairs of residues. For the dashed edges, only a single pair was identified.

**Table 1:** Key residues in the structural reorganisation of ErCry4 following photo-activation. The table summarizes hub residues with a degree larger than 12 and the 20 residues of highest betweenness centrality.

| Role | Residues |
| --- | --- |
| Hubs | Phe55, Leu57, Gln58, Ser59, Leu60, Asp62, Asn66, Leu67, Leu70, Thr83, Val84, Ile89, Lys108, Glu111, Ala112, Ile114, Gln115, Cys116, Leu117, Gly118, Phe123, Trp290, Phe293, Phe294, Val297, Ala298, Phe304, Asp339, Ala340, Thr343, Ile390, Gly393, Met396, Ile442, Gln453 |
| High betweenness | Glu12, Tyr35, Asp100, Glu102, Met103, Gly210, Leu219, Ser250, Tyr252, Arg261, Leu267, Thr333, Ala340, Leu345, Leu383, Asp385, Asp387, Tyr388, Met396, FAD (ribityl) |

**Table 2.**
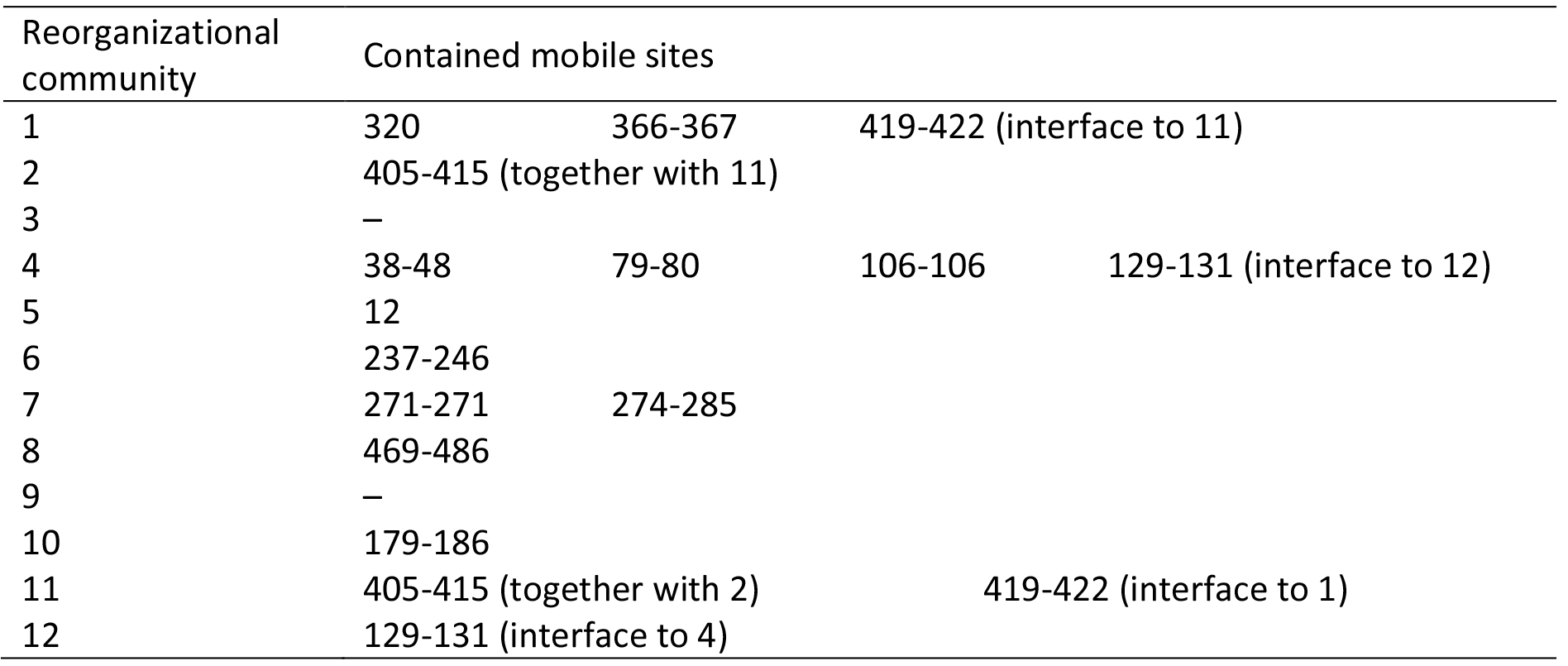
Distribution of mobile sites over reorganizational communities.

### Contact maps vs. dynamically correlated adjacency networks

With a description of the reorganizational dynamics of ErCry4 based on the correlated motion of adjacent segments established, the question arises if conventional contact networks can provide comparable insights. We here argue that this is not the case, because the mere adjacency of two residues does not necessarily imply an efficient reorganisation pathway. Only if spatially close sites undergo correlated displacements, can a functional relationship be inferred. Here, the presence of non-productive links in the contact network becomes obvious in the face of the markedly larger sparsity of the correlated displacement graphs, which eventually gives rise to altered reorganisation pathways. This is, for example, reflected in the distance of the mobile sites from the isoalloxazine subunit, F: While the contact network predicts that the site comprising residues 237 - 246 and the C-terminal lid (residues 405 - 415) are relatively close to F, the dynamic displacement map reveals that they are as peripheral as the CTT (compare Figures S10 and S13 in the SI). Likewise, residues that are identified as important to the contact network by their high betweenness centrality (e.g. His357, W_A_, W_D_, Asn394) are found not to actually transmit reorganisations by the correlated displacement maps. This is particularly obvious for His357, for which an analysis based on the contact network is likely to (erroneously) suggest functional relevancy. In strong contrast, the dynamic correlation network divulges that the residue is actually uncoupled from the main reorganisation network as a consequence of it lacking correlated motion with adjacent residues. Likewise, the contact network analysis overestimates the role of W_D_ in releasing the CTT by placing it at a bottleneck position of the shortest reorganisation path. While analysis of the correlated displacements based on eq. (7) reveals a shortest path close to this prediction, it indicates that Asp367 instead of Wd is important. The analysis based on eq. (6) favours an alternative pathway involving the α-helix H12. Thus, in summary, a study based on contact alone might fail to elucidate the intricate reorganisation properties of proteins.

### Phosphorylation sites

It is important to stress that some of the mobile residues of ErCry4 are serine, thus providing a hint to phosphorylation sites that could be functionally coupled to the photo-activation. In order to explore this possibility, we have used NetPhos 3.1 [73–74] to predict the propensity of serines, threonines or tyrosines to phosphorylation by kinases. We find that 8 out of the 42 predicted phosphorylation sites are located within mobile sites (see Table 3). Among these, the site surrounding Ser45 is particularly noteworthy as it comprises three adjacent phosphorylation sites (Thr43, Ser44, Ser45) that undergo α-helical coiling in response to the photo-activation of ErCry4. This conformational change is encoded by the second largest principal component accounting for 23 % of the positional variance encountered for the combined ensemble of DS and RPD states and is associated with a large increase in the SASA of Thr43 (see Figure S16). Likewise, the contact graphs show that the potential phosphorylation sites Ser180 and Ser278 are on average more exposed in the photo-activated state than the DS (Figure S9). While currently unsupported by experimental evidence, we speculate that phosphorylation at these sites could have a function in modulating/deactivating the photo-transduction cascade. This could proceed in a manner similar to the deactivation of rhodopsin, which after photoexcitation undergoes conformational changes that allow its phosphorylation by rhodopsin kinase [75]. It is then capped by arrestin thereby preventing further activation of transducine.

**Table 3.** Potential phosphorylation sites in regions that undergo large rearrangements upon the photo-activation of ErCry4. The table lists mobile regions by their residue sequence number and reorganisation communities using the numeric labels used in Figure 7.

| Potential phosphorylation sites | Mobile site | Reorganisation community |
| --- | --- | --- |
| T43, S44, S45 | 38 - 48 | 4 |
| S130 | 129 - 131 | 12 |
| S180 | 179 - 186 | 10 |
| S278 | 274 - 285 | 7 |
| T420 | 419 - 422 | 11 |
| S475 | CTT | 8 |

## Conclusions

The present investigation utilizes three tools to unravel the governing principles underlying the photoactivation of ErCry4. First, we have identified groups of residues that undergo marked structural reorganisation based on changes in internal coordinates, i.e. the distance matrix, and characterized their interrelation by principal component analysis. Second, we have mapped the protein structure onto a graph representation facilitated by the adjacency of residues and used these contact networks to identify common motifs of the mobile sites such as a low degree, an above-average local clustering coefficient and a low centrality. Third, we have demonstrated that these contact networks, though popular, can only reveal an incomplete picture of the dynamics. This shortcoming can be overcome by an alternative graph representation that is induced by the correlation of displacements in addition to the usual spatial adjacency. The signalling is multifaceted and involves many pathways: The network efficiently transmits correlation over short to intermediate distances via nearest-neighbour interactions and rigid secondary structure elements. The process is facilitated by hubs, which are four times more abundant than in the contact networks of the DS and RPD state. Tightly knit communities of residues, i.e. reorganisation modules, are interconnected by a small number of important residues, whose betweenness centrality also exceeds that of central nodes of the contact network by a factor of 3.5. Our analysis raises the prospect that the reorganisation properties could be altered or even engineered by manipulating these key links. In any case, the network layout provides a foundation for efficient signal transduction that goes beyond alternative modes of reorganisation such as a single pathway model or the undirected, i.e. stochastic, search of a new free energy minimum. More specifically, the analysis has revealed that the most prominent reorganisation sites in ErCry4 are located at the CTT and CTT-coupled motifs. In addition, the loop region involving Thr43-Ser45 has been found to show an interesting transition upon photo-activation whereupon a solvent-exposed α-helix forms. This and many of the other mobile sites expose serines (see Figure 5 and Table 3) that are potential phosphorylation sites in the ErCry4 photo-activation process. This observation allows us to insinuate that the photo-transduction cascade could potentially be modulated/controlled by kinases.

As the protein structure of ErCry4 has not yet been resolved, our study is based on a homology model. Despite the high degree of conservation within the photolyase/cryptochrome family of proteins, our predictions will have to be backed by analogous studies on the crystal structures once these become available. Furthermore, here we focus on a radical pair state that is formed in a photo-induced electron transfer reaction from the dark-state protein containing the FAD cofactor in the fully oxidized form. It is currently unknown and intensely debated whether this state could form the basis of a magnetic compass [10, 12, 25, 35, 43, 76–78]. Some evidence in favour of the alternative radical pair resulting from the re-oxidation of the fully reduced cryptochrome by e.g. oxygen has recently been found [14, 22, 36–42]. Studies of the signalling of this and related states of ErCry4 are currently ongoing. However, based on experimental findings for the analogous activation process in DmCry, we anticipate that the signalling behaviour will be prominently mediated by the cofactor and widely indifferent to the oxidation state of W_D_ [44]. The toolchain established here will prove useful in these follow-up studies.

Our approach has a bearing that goes beyond the photo-activation process in ErCry4 discussed here and is applicable to a multitude of signalling processes and allosteric adaption. In fact, it provides insights into a much deeper question: namely, does protein signalling/allostery involve dedicated pathways or follow a more holistic, concerted paradigm, for which the transition states and interactions are fleeting with only the initial and final state well defined. The ErCry4 studied here shows characteristics of both prototypes: concerted reorganisation in tightly coupled reorganisation modules but still remnants of pathways in the form of highly important residues at interface sites. Currently, we utilize extensive MD simulations to derive the correlation of displacements of residues. In doing so, we utilize graph representations to summarize the information contained in the trajectories. However, as the dynamically correlated networks are based on the concept of local correlation, this raises the prospect of deriving the graph representations from short-time MD simulations or alternative tools such as normal mode analysis or elastic network models [79]. In this way, the approach could provide predictive instead of descriptive power.

## Author contributions

D.R.K. and I.A.S. conceived and designed the study. D.R.K. and C.N. carried out the simulations and analysed the data. All authors discussed the results and contributed to the final manuscript.

## Acknowledgements

This project made use of time on ARCHER granted via the UK High-End Computing Consortium for Biomolecular Simulation, HECBioSim (www.hecbiosim.ac.uk), supported by EPSRC (grant no. EP/L000253/1). The authors would like to thank The Royal Society (RG170378), the EPSRC (grant no. EP/R021058/1) and the Lundbeck Foundation, Danish Councils for Independent Research and the Russian Science Foundation (Grant No. 17-72-2021) for financial support. Computational resources for the simulations were furthermore provided by the DeiC National HPC Center, SDU. DRK is thankful NVIDIA for supporting this research through their GPU Grant Program. This work used the UK Research Data Facility (http://www.archer.ac.uk/documentation/rdf-guide). Example input, parameter and output files for the NAMD simulations underlying this study are openly available from the University of Exeter’s institutional repository at: https://doi.org/10.24378/exe.363. This dataset includes a 100 ns trajectory of the RPD-state in the dcd-format (protein and FAD).

## References

1. Wiltschko, R.; Wiltschko, W., Magnetic Orientation in Animals. Springer: Berlin, New York, 1995; p 297 p.

2. Wiltschko, W., Über den Einfluss statischer Magnetfelder auf die Zugorienteirung der Rotkehlchen (Erithacus rubecula). Z. Tierpsychol. 1968, 25 (5), 537–58.

3. Wiltschko, W.; Merkel, F., Orientierung zugunruhiger Rotkehlchen im statischen Magnetfeld. Verh. Dtsch. Zool. Ges. 1966, 59, 362–367.

4. Wiltschko, W.; Munro, U.; Ford, H.; Wiltschko, R., Magnetic-inclination compass-a basis for the migratory orientation of birds in the northern and southern-hemisphere. Experientia 1993, 49 (2), 167–170.

5. Schulten, K.; Swenberg, C. E.; Weller, A., A biomagnetic sensory mechanism based on magnetic field modulated coherent electron spin motion. Z. Phys. Chem. 1978, 111 (1), 1–5.

6. Rodgers, C. T.; Hore, P. J., Chemical magnetoreception in birds: The radical pair mechanism. Proc. Natl. Acad. Sci. U. S. A. 2009, 106 (2), 353–360.

7. Ritz, T.; Adem, S.; Schulten, K., A model for photoreceptor-based magnetoreception in birds. Biophys. J. 2000, 78 (2), 707–718.

8. Chaves, I.; Pokorny,R.; Byrdin, M.; Hoang, N.; Ritz, T.; Brettel, K.; Essen, L. O.; van der Horst, G. T.; Batschauer, A.; Ahmad, M., The cryptochromes: blue light photoreceptors in plants and animals. Annu. Rev. Plant Biol. 2011, 62, 335–64.

9. Maeda, K.; Robinson, A. J.; Henbest, K. B.; Hogben, H. J.; Biskup, T.; Ahmad, M.; Schleicher, E.; Weber, S.; Timmel, C. R.; Hore, P. J., Magnetically sensitive light-induced reactions in cryptochrome are consistent with its proposed role as a magnetoreceptor. Proc. Natl. Acad. Sci. U. S. A. 2012, 109 (13), 4774–4779.

10. Lee, A. A.; Lau, J. C. S.; Hogben, H. J.; Biskup, T.; Kattnig, D. R.; Hore, P. J., Alternative radical pairs for cryptochrome-based magnetoreception. J. R. Soc., Interface 2014, 11 (95), 20131063.

11. Engels, S.; Schneider, N. L.; Lefeldt, N.; Hein, C. M.; Zapka, M.; Michalik, A.; Elbers, D.; Kittel, A.; Hore, P. J.; Mouritsen, H., Anthropogenic electromagnetic noise disrupts magnetic compass orientation in a migratory bird. Nature 2014, 509 (7500), 353–356.

12. Nielsen, C.; Kattnig, D. R.; Sjulstok, E.; Hore, P. J.; Solov’yov, I. A., Ascorbic acid may not be involved in cryptochrome-based magnetoreception. J. R. Soc., Interface 2017, 14 (137).

13. Kattnig, D. R.; Evans, E. W.; Dejean, V.; Dodson, C. A.; Wallace, M. I.; Mackenzie, S. R.; Timmel, C. R.; Hore, P. J., Chemical amplification of magnetic field effects relevant to avian magnetoreception. Nat. Chem. 2016, 8 (4), 384–391.

14. Wiltschko, R.; Ahmad, M.; Nießner, C.; Gehring, D.; Wiltschko, W., Light-dependent magnetoreception in birds: The crucial step occurs in the dark. J. R. Soc., Interface 2016, 13 (118), 20151010.

15. Schwarze, S.; Schneider, N. L.; Reichl, T.; Dreyer, D.; Lefeldt, N.; Engels, S.; Baker, N.; Hore, P. J.; Mouritsen, H., Weak broadband electromagnetic fields are more disruptive to magnetic compass orientation in a night-migratory songbird (Erithacus rubecula) than strong narrow-band fields. Front. Behav. Neurosci. 2016, 10, 55.

16. Günther, A.; Einwich, A.; Sjulstok, E.; Feederle, R.; Bolte, P.; Koch, K. W.; Solov’yov, I. A.; Mouritsen, H., Double-Cone Localization and Seasonal Expression Pattern Suggest a Role in Magnetoreception for European Robin Cryptochrome 4. Curr. Biol. 2018, 28 (2), 211–223.

17. Pinzon-Rodriguez, A.; Bensch, S.; Muheim, R., Expression patterns of cryptochrome genes in avian retina suggest involvement of Cry4 in light-dependent magnetoreception. J. R. Soc., Interface 2018, 15 (140).

18. Solov’yov, I. A.; Schulten, K., Reaction Kinetics and Mechanism of Magnetic Field Effects in Cryptochrome. J. Phys. Chem. B 2012, 116 (3), 1089–1099.

19. Bazalova, O.; Kvicalova, M.; Valkova, T.; Slaby, P.; Bartos, P.; Netusil, R.; Tomanova, K.; Braeunig, P.; Lee, H. J.; Sauman, I.; Damulewicz, M.; Provaznik, J.; Pokorny, R.; Dolezel, D.; Vacha, M., Cryptochrome 2 mediates directional magnetoreception in cockroaches. Proc. Natl. Acad. Sci. U. S. A. 2016, 113 (6), 1660–1665.

20. Bolte, P.; Bleibaum, F.; Einwich, A.; Gunther, A.; Liedvogel, M.; Heyers, D.; Depping, A.; Wohlbrand, L.; Rabus, R.; Janssen-Bienhold, U.; Mouritsen, H., Localisation of the Putative Magnetoreceptive Protein Cryptochrome 1b in the Retinae of Migratory Birds and Homing Pigeons. Plos One 2016, 11 (3).

21. Weber, S.; Biskup, T.; Okafuji, A.; Marino, A. R.; Berthold, T.; Link, G.; Hitomi, K.; Getzoff, E. D.; Schleicher, E.; Norris, J. R., Origin of light-induced spin-correlated radical pairs in cryptochrome. J. Phys. Chem. B 2010, 114 (45), 14745–14754.

22. Wiltschko, R.; Gehring, D.; Denzau, S.; Nießner, C.; Wiltschko, W., Magnetoreception in birds: II. Behavioural experiments concerning the cryptochrome cycle. J. Exp. Biol. 2014, 217 (Pt 23), 4225–8.

23. Wiltschko, R.; Wiltschko, W., Sensing magnetic directions in birds: radical pair processes involving cryptochrome. Biosensors 2014, 4 (3), 221–42.

24. Sheppard, D. M. W.; Li, J.; Henbest, K. B.; Neil, S. R. T.; Maeda, K.; Storey, J.; Schleicher, E.; Biskup, T.; Rodriguez, R.; Weber, S.; Hore, P. J.; Timmel, C. R.; Mackenzie, S. R., Millitesla magnetic field effects on the photocycle of an animal cryptochrome. Sci. Rep. 2017, 7, 42228.

25. Hore, P. J.; Mouritsen, H., The radical-pair mechanism of magnetoreception. Annu. Rev. Biophys. 2016, 45, 299–344.

26. Muheim, R.; Boström, J.; Åkesson, S.; Liedvogel, M., Sensory mechanisms of animal orientation and navigation. In Animal Movement Across Scales, Åkesson, L.-A. H. S., Ed. Oxford University Press: 2014; pp 179–194.

27. Liedvogel, M.; Maeda, K.; Henbest, K.; Schleicher, E.; Simon, T.; Timmel, C. R.; Hore, P. J.; Mouritsen, H., Chemical magnetoreception: Bird cryptochrome 1a is excited by blue light and forms long-lived radical-pairs. PLoS One 2007, 2 (10), e1106.

28. Mitsui, H.; Maeda, T.; Yamaguchi, C.; Tsuji, Y.; Watari, R.; Kubo, Y.; Okano, K.; Okano, T., Overexpression in Yeast, Photocycle, and in Vitro Structural Change of an Avian Putative Magnetoreceptor Cryptochrome4. Biochem. 2015, 54 (10), 1908–1917.

29. Mouritsen, H.; Janssen-Bienhold, U.; Liedvogel, M.; Feenders, G.; Stalleicken, J.; Dirks, P.; Weiler, R., Cryptochromes and neuronal-activity markers colocalize in the retina of migratory birds during magnetic orientation. Proc. Natl. Acad. Sci. U. S. A. 2004, 101 (39), 14294–14299.

30. Nießner, C.; Denzau, S.; Gross, J. C.; Peichl, L.; Bischof, H. J.; Fleissner, G.; Wiltschko, W.; Wiltschko, R., Avian Ultraviolet/Violet Cones Identified as Probable Magnetoreceptors. Plos One 2011, 6 (5).

31. Nießner, C.; Gross, J. C.; Denzau, S.; Peichl, L.; Fleissner, G.; Wiltschko, W.; Wiltschko, R., Seasonally Changing Cryptochrome 1b Expression in the Retinal Ganglion Cells of a Migrating Passerine Bird. Plos One 2016, 11 (3).

32. Kutta, R. J.; Archipowa, N.; Johannissen, L. O.; Jones, A. R.; Scrutton, N. S., Vertebrate Cryptochromes are Vestigial Flavoproteins. Sci. Rep. 2017, 7.

33. Ozturk, N.; Selby, C. P.; Song, S. H.; Ye, R.; Tan, C.; Kao, Y. T.; Zhong, D. P.; Sancar, A., Comparative Photochemistry of Animal Type 1 and Type 4 Cryptochromes. Biochem. 2009, 48 (36), 8585–8593.

34. Watari, R.; Yamaguchi, C.; Zemba, W.; Kubo, Y.; Okano, K.; Okano, T., Light-dependent Structural Change of Chicken Retinal Cryptochrome4. J. Biol. Chem. 2012, 287 (51), 42634–42641.

35. Hiscock, H. G.; Worster, S.; Kattnig, D. R.; Steers, C.; Jin, Y.; Manolopoulos, D. E.; Mouritsen, H.; Hore, P. J., The quantum needle of the avian magnetic compass. Proc. Natl. Acad. Sci. U. S. A. 2016, 113 (17), 4634–4639.

36. Wiltschko, R.; Stapput, K.; Thalau, P.; Wiltschko, W., Directional orientation of birds by the magnetic field under different light conditions. J. R. Soc., Interface 2010, 7 Suppl 2, S163–77.

37. Nießner, C.; Denzau, S.; Stapput, K.; Ahmad, M.; Peichl, L.; Wiltschko, W.; Wiltschko, R., Magnetoreception: activated cryptochrome 1a concurs with magnetic orientation in birds. J. R. Soc., Interface 2013, 10 (88), 20130638.

38. Nießner, C.; Denzau, S.; Peichl, L.; Wiltschko, W.; Wiltschko, R., Magnetoreception in birds:I. Immunohistochemical studies concerning the cryptochrome cycle. J. Exp. Biol. 2014, 217 (Pt 23), 4221–4.

39. Ritz, T.; Wiltschko, R.; Hore, P. J.; Rodgers, C. T.; Stapput, K.; Thalau, P.; Timmel, C. R.; Wiltschko, W., Magnetic compass of birds is based on a molecule with optimal directional sensitivity. Biophys. J. 2009, 96 (8), 3451–3457.

40. Müller, P.; Ahmad, M., Light-activated cryptochrome reacts with molecular oxygen to form a flavin-superoxide radical pair consistent with magnetoreception. J. Biol. Chem. 2011, 286 (24), 21033–40.

41. Solov’yov, I. A.; Schulten, K., Magnetoreception through Cryptochrome May Involve Superoxide. Biophys. J. 2009, 96 (12), 4804–4813.

42. Kattnig, D. R., Radical-Pair-Based Magnetoreception Amplified by Radical Scavenging: Resilience to Spin Relaxation. J. Phys. Chem. B 2017, 121 (44), 10215–10227.

43. Kattnig, D. R.; Sowa, J. K.; Solov’yov, I. A.; Hore, P. J., Electron spin relaxation can enhance the performance of a cryptochrome-based magnetic compass sensor. New J. Phys. 2016, 18, 063007.

44. Vaidya, A. T.; Top, D.; Manahan, C. C.; Tokuda, J. M.; Zhang, S.; Pollack, L.; Young, M. W.; Crane, B. R., Flavin reduction activates Drosophila cryptochrome. Proc. Natl. Acad. Sci. U. S. A. 2013, 110 (51), 20455–20460.

45. Ganguly, A.; Manahan, C. C.; Top, D.; Yee, E. F.; Lin, C. F.; Young, M. W.; Thiel, W.; Crane, B. R., Changes in active site histidine hydrogen bonding trigger cryptochrome activation. Proc. Natl. Acad. Sci. U. S. A. 2016, 113 (36), 10073–10078.

46. Thöing, C.; Oldemeyer, S.; Kottke, T., Microsecond Deprotonation of Aspartic Acid and Response of the alpha/beta Subdomain Precede C-Terminal Signaling in the Blue Light Sensor Plant Cryptochrome. J. Am. Chem. Soc. 2015, 137 (18), 5990–5999.

47. Hense, A.; Herman, E.; Oldemeyer, S.; Kottke, T., Proton Transfer to Flavin Stabilizes the Signaling State of the Blue Light Receptor Plant Cryptochrome. J. Biol. Chem. 2015, 290 (3), 1743–1751.

48. He, S. B.; Wang, W. X.; Zhang, J. Y.; Xu, F.; Lian, H. L.; Li, L.; Yang, H. Q., The CNT1 Domain of Arabidopsis CRY1 Alone Is Sufficient to Mediate Blue Light Inhibition of Hypocotyl Elongation. Mol. Plant 2015, 8 (5), 822–825.

49. Kondoh, M.; Shiraishi, C.; Muller, P.; Ahmad, M.; Hitomi, K.; Getzoff, E. D.; Terazima, M., Light-Induced Conformational Changes in Full-Length Arabidopsis thaliana Cryptochrome. J. Mol. Biol. 2011, 413 (1), 128–137.

50. Ahmad, M., Photocycle and signaling mechanisms of plant cryptochromes. Curr. Opin. Plant Biol. 2016, 33, 108–115.

51. Hünenberger, P. H.; Mark, A. E.; Vangunsteren, W. F., Fluctuation and Cross-Correlation Analysis of Protein Motions Observed in Nanosecond Molecular-Dynamics Simulations. J. Mol. Biol. 1995, 252 (4), 492–503.

52. Karplus, M.; Ichiye, T., Fluctuation and cross correlation analysis of protein motions observed in nanosecond molecular dynamics simulations. J. Mol. Biol. 1996, 263 (2), 120–122.

53. Ichiye, T.; Karplus, M., Collective Motions in Proteins-a Covariance Analysis of Atomic Fluctuations in Molecular-Dynamics and Normal Mode Simulations. PROTEINS 1991, 11 (3), 205–217.

54. Di Paola, L.; De Ruvo, M.; Paci, P.; Santoni, D.; Giuliani, A., Protein Contact Networks: An Emerging Paradigm in Chemistry. Chem. Rev. 2013, 113 (3), 1598–1613.

55. Kasahara, K.; Fukuda, I.; Nakamura, H., A Novel Approach of Dynamic Cross Correlation Analysis on Molecular Dynamics Simulations and Its Application to Ets1 Dimer-DNA Complex. Plos One 2014, 9 (11).

56. Yoon, H. J.; Lee, S.; Park, S. J.; Wu, S., Network approach of the conformational change of c-Src, a tyrosine kinase, by molecular dynamics simulation. Sci Rep 2018, 8 (1), 5673.

57. Phillips, J. C.; Braun, R.; Wang, W.; Gumbart, J.; Tajkhorshid, E.; Villa, E.; Chipot, C.; Skeel, R. D.; Kale, L.; Schulten, K., Scalable molecular dynamics with NAMD. J. Comput. Chem. 2005, 26 (16), 1781–1802.

58. Best, R. B.; Zhu, X.; Shim, J.; Lopes, P. E. M.; Mittal, J.; Feig, M.; MacKerell, A. D., Optimization of the additive CHARMM all-atom protein force field targeting improved sampling of the backbone ϕ ψ and side-chain χ_1_ and χ_2_ dihedral angles. J. Chem. Theo. Comp. 2012, 8 (9), 3257–3273.

59. MacKerell, A. D.; Feig, M.; Brooks, C. L., Extending the treatment of backbone energetics in protein force fields: Limitations of gas-phase quantum mechanics in reproducing protein conformational distributions in molecular dynamics simulations. J. Comput. Chem. 2004, 25 (11), 1400–1415.

60. MacKerell, A. D.; Bashford, D.; Bellott, M.; Dunbrack, R. L.; Evanseck, J. D.; Field, M. J.; Fischer, S.; Gao, J.; Guo, H.; Ha, S.; Joseph-McCarthy, D.; Kuchnir, L.; Kuczera, K.; Lau, F. T. K.; Mattos, C.; Michnick, S.; Ngo, T.; Nguyen, D. T.; Prodhom, B.; Reiher, W. E.; Roux, B.; Schlenkrich, M.; Smith, J. C.; Stote, R.; Straub, J.; Watanabe, M.; Wiorkiewicz-Kuczera, J.; Yin, D.; Karplus, M., All-atom empirical potential for molecular modeling and dynamics studies of proteins. J. Phys. Chem. B 1998, 102 (18), 3586–3616.

61. Lüdemann, G.; Solov’yov, I. A.; Kubar, T.; Elstner, M., Solvent driving force ensures fast formation of a persistent and well-separated radical pair in plant cryptochrome. J. Am. Chem. Soc. 2015, 137 (3), 1147–1156.

62. Solov’yov, I. A.; Domratcheva, T.; Schulten, K., Separation of photo-induced radical pair in cryptochrome to a functionally critical distance. Sci. Rep. 2014, 4, 3845

63. Sjulstok, E.; Olsen, J. M. H.; Solov’yov, I. A., Quantifying electron transfer reactions in biological systems: what interactions play the major role? Sci. Rep. 2015, 5, 18446.

64. Gaci, O., A topological description of hubs in amino Acid interaction networks. Adv Bioinformatics 2010, 257512.

65. Lienin, S. F.; Bruschweiler, R., Characterization of collective and anisotropic reorientational protein dynamics. Phys. Rev. Lett. 2000, 84 (23), 5439–5442.

66. Frishman, D.; Argos, P., Knowledge-based protein secondary structure assignment. PROTEINS 1995, 23 (4), 566–579.

67. David, C. C.; Jacobs, D. J., Principal component analysis: a method for determining the essential dynamics of proteins. Methods Mol. Biol. 2014, 1084, 193–226.

68. Wolfram Research, I., Mathematica. Wolfram Research, Inc.: Champaign, Illinois, 2018.

69. Solov’yov, I. A.; Domratcheva, T.; Shahi, A. R. M.; Schulten, K., Decrypting Cryptochrome: Revealing the Molecular Identity of the Photoactivation Reaction. J. Am. Chem. Soc. 2012, 134 (43), 18046–18052.

70. Kottke, T.; Batschauer, A.; Ahmad, M.; Heberle, J., Blue-light-induced changes in Arabidopsis cryptochrome 1 probed by FTIR difference spectroscopy. Biochem. 2006, 45 (8), 2472–2479.

71. Langenbacher, T.; Immeln, D.; Dick, B.; Kottke, T., Microsecond Light-induced Proton Transfer to Flavin in the Blue Light Sensor Plant Cryptochrome. J. Am. Chem. Soc. 2009, 131 (40), 14274–14280.

72. Brettel, K.; Byrdin, M., Reaction mechanisms of DNA photolyase. Curr. Opin. Struct. Biol. 2010, 20 (6), 693–701.

73. Blom, N.; Sicheritz-Ponten, T.; Gupta, R.; Gammeltoft, S.; Brunak, S., Prediction of post-translational glycosylation and phosphorylation of proteins from the amino acid sequence. Proteomics 2004, 4 (6), 1633–1649.

74. Blom, N.; Gammeltoft, S.; Brunak, S., Sequence and structure-based prediction of eukaryotic protein phosphorylation sites. J. Mol. Biol. 1999, 294 (5), 1351–1362.

75. Koch, K. W.; Dell’Orco, D., Protein and Signaling Networks in Vertebrate Photoreceptor Cells. Front. Mol. Neurosci. 2015, 8.

76. Kattnig, D. R.; Hore, P. J., The sensitivity of a radical pair compass magnetoreceptor can be significantly amplified by radical scavengers. Sci. Rep. 2017, 7 (1), 11640.

77. Kattnig, D. R.; Solov’yov, I. A.; Hore, P. J., Electron spin relaxation in cryptochrome-based magnetoreception. Phys. Chem. Chem. Phys. 2016, 18 (18), 12443–12456.

78. Pedersen, J. B.; Nielsen, C.; Solov’yov, I. A., Multiscale description of avian migration: from chemical compass to behaviour modeling. Sci. Rep. 2016, 6, 36709.

79. Bahar, I.; Lezon, T. R.; Bakan, A.; Shrivastava, I. H., Normal Mode Analysis of Biomolecular Structures: Functional Mechanisms of Membrane Proteins. Chem. Rev. 2010, 110 (3), 1463–1497.

